# A Combined Spectroscopic and Molecular Modeling Study on Structure-Function-Dynamics under Chemical Modification: Alpha-Chymotrypsin with Formalin Preservative

**DOI:** 10.1101/2021.07.24.453635

**Authors:** Pritam Biswas, Aniruddha Adhikari, Uttam Pal, Susmita Mondal, Dipanjan Mukherjee, Ria Ghosh, Tanusri Saha-Dasgupta, Sudeshna Shyam Chowdhury, Ranjan Das, Samir Kumar Pal

**Affiliations:** Department of Microbiology, St. Xavier’s College, 30, Mother Teresa Sarani, Kolkata 700016, India; Department of Chemical, Biological and Macromolecular Sciences, S. N. Bose National Centre for Basic Sciences, Block JD, Sector III, Salt Lake, Kolkata 700106, India; Technical Research Centre, S. N. Bose National Centre for Basic Sciences, Block JD, Sector III, Salt Lake, Kolkata 700106, India; Department of Condensed Matter Physics and Material Sciences, S. N. Bose National Centre for Basic Sciences, Block JD, Sector III, Salt Lake, Kolkata 700106, India; Department of Chemistry, West Bengal state University, Barasat, Kolkata 700126, India

**Keywords:** amino acid modification, chymotrypsin, formalin, cross-linking, spectroscopy, molecular docking

## Abstract

Enzyme conformations can be altered via modification of its amino acid residues, side chains and large-scale domain modifications, which are closely linked to its function. Herein, we have addressed the role of residue modification in catalytic activity and molecular recognition of an enzyme alpha-chymotrypsin (CHT) in presence of covalent cross-linker formalin. Optical spectroscopy studies exhibit reduced catalytic activity of the enzyme with increased formalin concentration. Polarization gated anisotropy studies of a fluorophore 8-anilino-1- napthelenesulfonic acid (ANS) in CHT show a dip rise pattern in presence of formalin which is consistent with the generation of multiple ANS binding sites in the enzyme owing to modifications of its local amino acid residues. Molecular docking study on minimal local residue modifications in CHT reveals formation of a stable enzyme-substrate complex even with the serine-histidine cross-linked enzyme which prohibits product formation giving rise to reduced catalytic activity.

## Introduction

Covalent modifications introduce addition or removal of groups from the amino acid residues of a protein altering its structure and function. Formalin is known to react with a plethora of functional groups in small peptides and other substrates to generate covalently modified products [1, 2] which can include both intramolecular and intermolecular cross-linked species. From a chemical standpoint, formalin can react with biological nucleophiles such as protein and DNA and facilitates *in vitro* formation of intra-strand and DNA-protein cross links [2–4]. Formalin related reactions in cells are likely responsible for its toxic/carcinogenic effects, however the precise chemistry of such reactions and their cellular prevalence is still not clearly defined [5]. Pioneering works by McGhee and Von Hippel [6] and a recent study by Kamps et al. [7] on the reactions of nucleosides/nucleotides and amino acids with formalin have envisaged to understand the chemistry of such reactions.

To have a better understanding how formalin influence health and diseases, it is important to investigate the correlation between formalin induced structural modification and function of biological macromolecules. Previous studies on interaction of formalin with biological macromolecules were restricted to model peptides and other proteins [4, 8–10]. These studies, however, revealed very little information on residue modification, and its correlation to protein function and its molecular recognition. In the present study, residue modification of a common digestive enzyme α-chymotrypsin (CHT; EC 3.4.21.1) by formalin was correlated with its catalytic activity and molecular recognition of a ligand (ANS) using steady state and pico-second resolved fluorescence spectroscopy, circular dichroism (CD) in conjunction with molecular docking studies.

## Materials and methods

### Materials

α-Chymotrypsin (CHT) (*Bostaurus*) (≥ 40U/mg), Ala-Ala-Phe-7-amido-4-methylcoumarin (AMC) (≥98%), 8-Anilino-1-napthelenesulfonic acid (ANS) (≥97%) were purchased from Sigma (Saint Louis, USA). α-Chymotrypsin, AMC and ANS solutions were prepared in phosphate buffer (10mM, pH 7.0) using water from Millipore system.

### Sample preparation

The ANS-CHT complex was prepared by mixing ANS (0.5 μM) with CHT (5 μM) in phosphate buffer (pH 7.0). The mix was stirred continuously for 5 h at 4°C, which was filtered extensively to remove the free probe [11].

### Circular Dichroism (CD) Spectroscopy

Circular dichroism (CD) spectroscopy analysis of CHT was conducted with 2 μM CHT solution incubated with different concentrations of formalin (1%, 2%, 4%) using a quartz cuvette of 1 mm path length in a JASCO 815 spectrometer. The deconvolution of the CD signals into relevant secondary structure was done by CDNN software [12].

### Dynamic light scattering (DLS)

Dynamic light scattering (DLS) measurements was done with 1.5 μM CHT solution incubated with different concentrations of formalin using a Nano S Malvern instrument employing a 4 mW He-Ne laser (λ_ex_ = 632 nm).

### Enzyme Activity Assay

Enzymatic activity of CHT was conducted with 0.5 μM of CHT at different formalin concentrations (1%, 2%, 4%). The substrate Ala-Ala-Phe-AMC (λ_max_ = 325 nm) was cleaved to produce 7-amido-4-methyl-coumarin (λ_max_ = 370 nm), the absorbance of which was monitored in Shimadzu Model UV-2600 spectrophotometer.

### Time correlated single photon counting measurement

Picosecond-resolved fluorescence spectroscopic study were carried out using a commercial time correlated single photon counting (TCSPC) from Edinburgh instruments. The picosecond excitation pulses from the picoquant diode laser were used at 375 nm for the excitation of ANS with an instrument response function (IRF) of 80 ps at different formalin concentration. Fluorescence photons were detected by a microchannel-plate-photomultiplier tube (MCP-PMT; Hamamatsu Photonics, Kyoto, Japan). The time-resolved instrument provided a time resolution of 20 ps, which is one-fourth of the instrument response (IRF) for the detection of time constants after de-convolution of the IRF. For all decays, the emission polarizer was set at 55° (magic angle) with respect to the polarization axis of the emission beam. The observed fluorescence transients were fitted using a nonlinear least-squares fitting procedure. The details of which can be found elsewhere [13, 14].

### Polarization-gated anisotropy measurements

For the anisotropy measurements (r(*t*)), the emission polarizer was adjusted to be parallel and perpendicular to that of the excitation. Time-resolved anisotropy, r(*t*), was determined from the following equation:

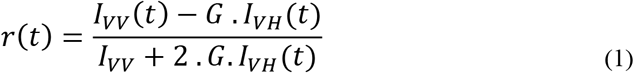

where *I*_VV_(*t*) and *I*_VH_(*t*) are parallel and perpendicular polarized fluorescence decays of the dye, respectively, recorded using a vertically polarized excitation light. *G* is an instrument grating factor, which is an instrument and wavelength dependent correction factor to compensate for the polarization bias of the detection system and its magnitude was obtained by a long tail matching technique [15, 16].

### Solvent accessible surface area (SASA)

Solvent accessible surface area (SASA) is a measure of formation of contacts between the atoms on the surface of the protein and the solvent molecules. Thus, solvent accessibility provides a better insight on the amino acid residues of the enzyme (CHT) vulnerable to formalin induced modification. The validation and selection of the appropriate protein data bank (PDB) structure of CHT was made according to Chakraborty et al.[17]. Solvent accessible surface area (SASA) of CHT (PDB ID: 1CGJ) is computed using the PyMOL software (Educational version). The catalytic and residues of ANS binding site were identified as the prime targets for the SASA analysis.

### Residue modification and molecular docking

Molecular docking was performed to determine the catalytic and the ANS binding residues using AutoDock Vina [18]. Solvent exposed residues susceptible to formalin modification were modified followed by minimization of the modified residues using Schrodinger Maestro 2018-1 (Academic release). All covalent modifications were conducted according to Kamps et al.[7] Molecular docking studies were performed to study the influence of formalin induced covalent modification on substrate and molecular recognition of ANS by CHT. Furthermore, SASA analyses of the bound ANS were performed according to the methodology described in the previous section.

## Results and Discussion

### Structural analysis

α-chymotrypsin (CHT) is a proteolytic enzyme of the class serine protease (EC 3.4.21.1) which is associated with hydrolysis of peptide bonds in the mammalian digestive system. The catalytic triad of CHT is located in the hydrophobic S1 pocket of CHT [19], molecular docking studies of CHT-ANS indicates the presence of two different sites-one located at the S1 pocket (buried internal site) while the other at the cysteine-1-122 disulfide bond (exposed external site). Fig. 1 shows the 3D structure of CHT. Fig 1(a) highlights the catalytic triad comprising the residues histidine-57; aspartate-102 and serine-195. The site at cys-1-122 disulfide bond (Fig 1b) serves as one of the ANS binding sites and is located almost opposite to the catalytic site.

**Fig. 1.**
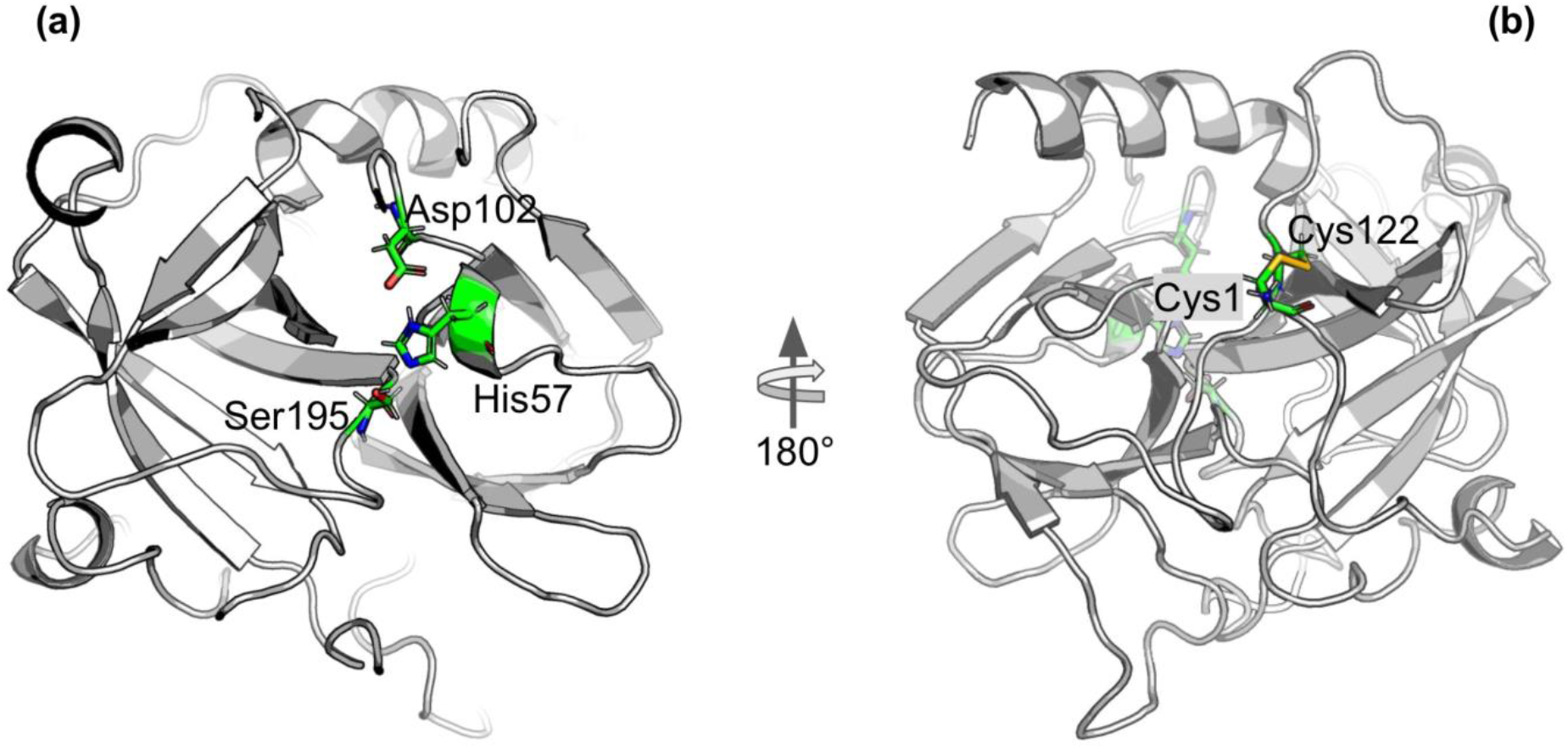
Cartoon representation of alpha-Chymotrypsin (CHT) [PDB ID:1CGJ]. (a) The coloured stick representation of the active site catalytic triad; showing the active site residues [His-57;Asp-102;Ser-195]. (b) At the N-terminal, the coloured stick representation [Cys-1; Cys-122] shows the molecular recognition site [External ANS binding site]. This site is located almost opposite to the catalytic site.

### Enzymatic activity assay

The catalytic activity of CHT was measured with different AMC (substrate) concentrations (5-40 μM) at varying formalin concentrations (1% to 4%). CHT belongs to the class of serine protease that catalyzes hydrolysis of AMC to 7-amido-4-methy-coumarin (Fig 2a & b). Fig 2c & d illustrate the effects of increasing formalin concentration on the product formation and the rate of product formation. A reduction in the rate of CHT catalyzed product formation was observed (Table 1) with increasing formalin concentration reflecting on reduced catalytic activity which likely originates from a possible modification of the substrate binding site.

**Fig. 2:**
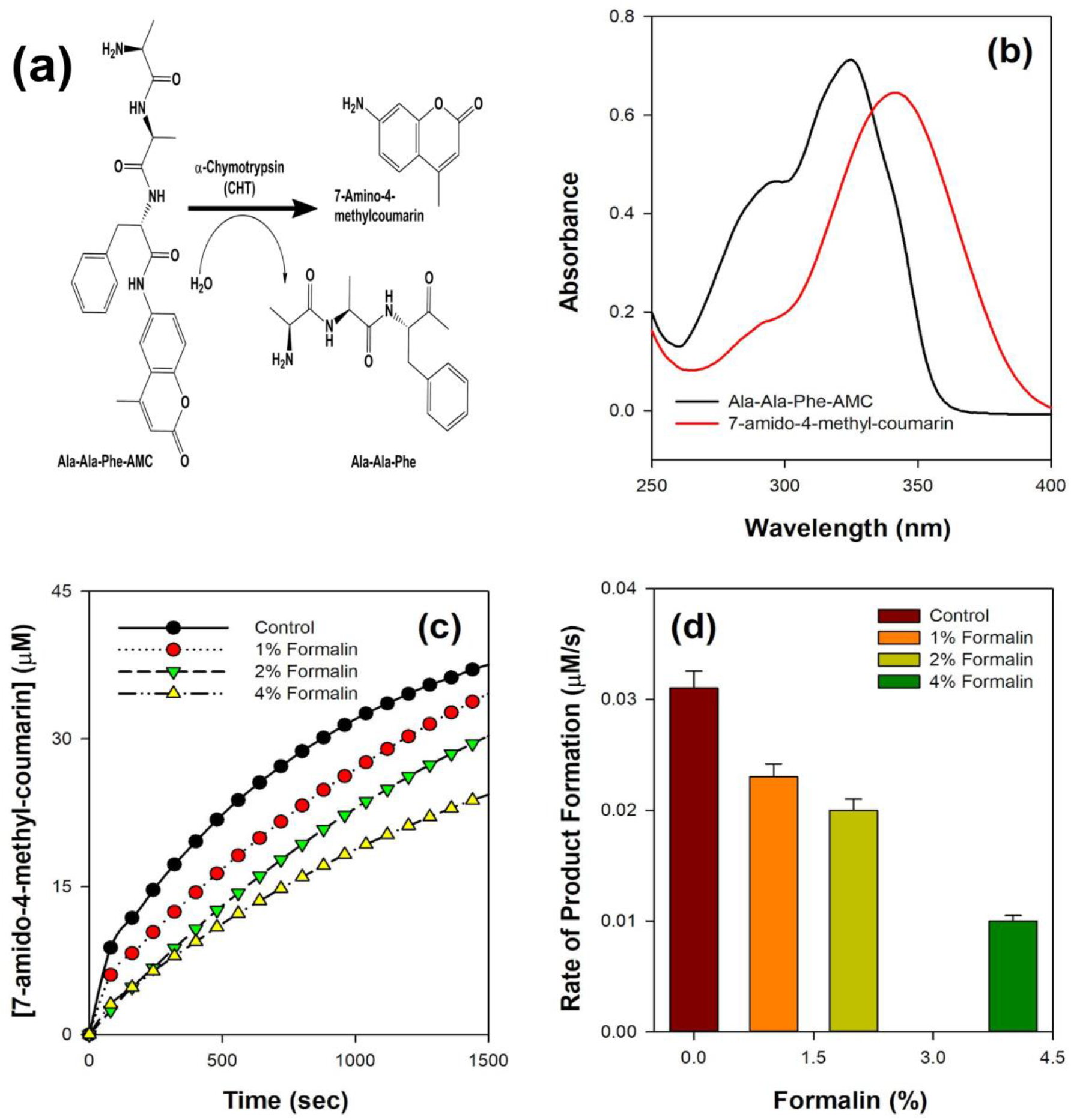
Schematic representation of the formalin treated enzyme catalysis. (a) Simple reaction outline catalysed by CHT; the substrate AMC hydrolysed to produce 7-amido-4-methyl-coumarin. (b) Absorbance spectra of the substrate AMC and product 7-amido-4-methyl-coumarin during catalysis. (c) Product formation at different representative formalin concentration; diminishing product formation with increased formalin concentration. (d) Rate of product formation at different representative formalin concentration; retarding rate of product formation consistent with decreasing product formation with increasing formalin concentration.

**Table 1.** Rate of product formation of alpha-Chymotrypsin at different formalin concentration.

| Formalin Concentration (%) | Rate of product formation ($\mu\text{M/s}$) |
| --- | --- |
| Control | $(0.031 \pm 1.55) \times 10^{-3}$ |
| 1 | $(0.023 \pm 1.15) \times 10^{-3}$ |
| 2 | $(0.020 \pm 1.0) \times 10^{-3}$ |
| 4 | $(0.010 \pm 5.0) \times 10^{-4}$ |

### Secondary and globular tertiary structure analysis

Close analysis of the 3D structure of CHT (Fig 1) shows that the enzyme consists of anti-parallel β-sheets which are highly distorted, forming very short irregular strands. Such conformation can lead to a shift in the negative band from the ideal β-sheet position (210-220 nm) towards 200 nm.

The effect of formalin induced modification on CHT was quantified by circular dichroism (CD) spectroscopy in the far UV region. The far UV spectra of CHT were characterized by a minimum at 202 nm with no positive band (Fig 3). Upon de-convolution, a 4% increase in anti-parallel β-sheet content was observed along with a nominal decrease in α-helix and insignificant changes in parallel β-sheet, turns and random coils (Table 2). These results indicate an insignificant change in the secondary structure of the enzyme reflecting on its structural stability.

**Fig. 3:**
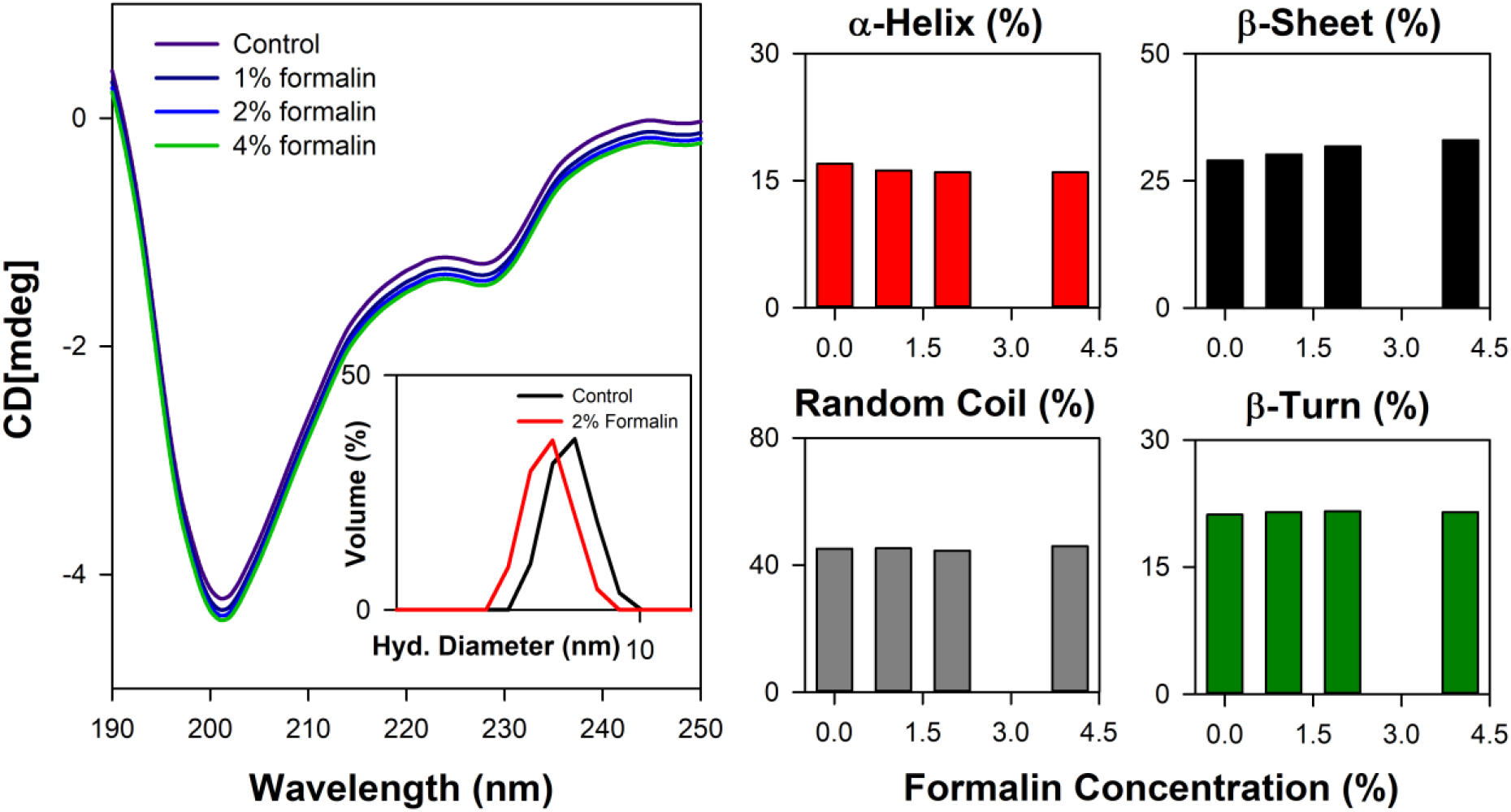
Far UV CD spectroscopy and DLS study of CHT. (a) The far UV CD spectra representing structural integrity of formalin treated CHT. The inset shows the DLS spectra of CHT at representative formalin concentration. (b) Percentage composition of secondary structure shown as bar plots. Analysis of secondary structure indicated structural stability within the experimental setup.

**Table 2.** Secondary structure analysis of alpha-Chymotrypsin in presence of formalin.

| Formalin conc (%) | % α-helix | % β-sheet | % random coil | % β-turn |
| --- | --- | --- | --- | --- |
| Control | 17.00 | 29.00 | 45.20 | 21.20 |
| 1 | 16.20 | 30.20 | 45.30 | 21.50 |
| 2 | 16.00 | 31.80 | 44.60 | 21.60 |
| 4 | 16.00 | 33.00 | 46.00 | 21.50 |

The effect of formalin on the globular tertiary structure of CHT was monitored by dynamic light scattering (DLS). The DLS study (Fig 3, inset) reveals the hydrodynamic diameter of CHT to be ~7 nm, which agrees with previous studies [20, 21]. Upon treatment with 4% formalin, a decrease in the hydrodynamic diameter of CHT from ~ 7 nm (control) to ~ 5 nm was noted (Table S1, supporting information file) indicating increased compactness and rigidity rather than structural unfolding of the enzyme over the range of formalin concentration used.

### Picosecond-resolved Fluorescence Studies

In water fluorescence of ANS decays rapidly with a time constant of 0.25 ns [22]. The fluorescence transients of ANS in CHT at different formalin concentrations (Fig. 4) are characterized by three-time constants (Table 3) indicating its binding to the enzyme. The faster component (τ_1_) of few hundreds of picoseconds (~ 0.25 ns) originates due to free ANS in water [22]. ANS bound to CHT reveals two different lifetimes (~ 2.1 ns and ~ 7.1 ns) which may be ascribed to two different binding sites in the enzyme; the shorter lifetime (τ_2_) corresponds to an external binding site partially exposed to water, whereas, the longer one (τ_3_) may be ascribed to an internal binding site located inside the enzyme which is screened from water. Upon increasing formalin content, the longer time constant remains unchanged whereas the shorter fluorescence lifetime (τ_2_) decreases to 1.4 ns (Table 3). This may be ascribed to structural modification in the external binding site leading to increased exposure of the probe (ANS) to water resulting in a shortening of the shorter fluorescence lifetime (τ_2_).

**Fig. 4:**
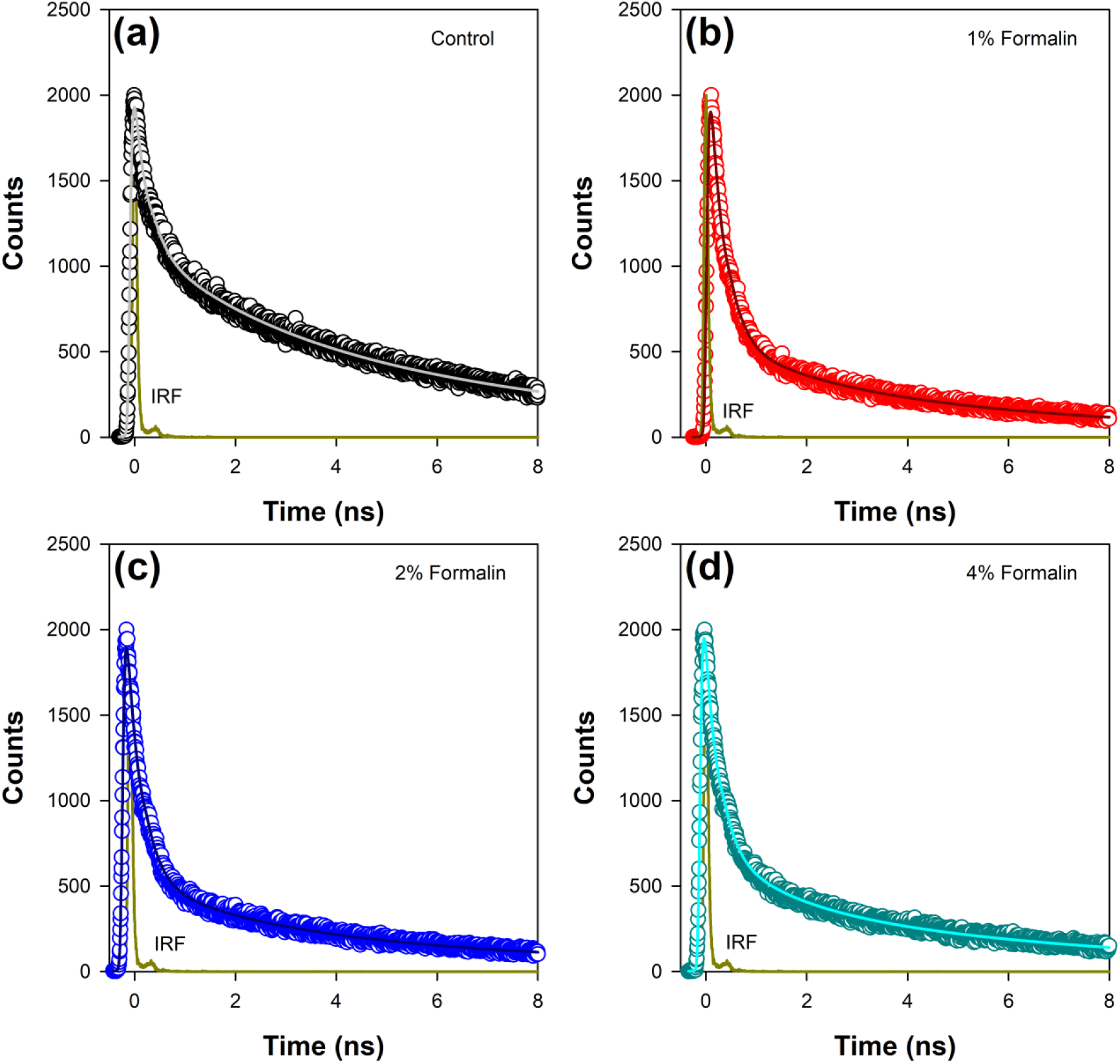
Time resolved fluorescence transients of ANS in (a) pure CHT (control) and at varying formalin concentrations (b-d).

**Table. 3:** Fluorescence lifetimes of ANS in α-Chymotrypsin in presence of formalin.

| Formalin conc. (%) | $\tau_1$ (ps) | $\tau_2$ (ps) | $\tau_3$ (ps) | $\tau_{\text{avg}}$ (ps) |
| --- | --- | --- | --- | --- |
| Control | 0.235 (55) | 2.11 (14) | 7.05 (31) | 2.71 |
| 1 | 0.259 (78) | 1.94 (10) | 7.21 (12) | 1.29 |
| 2 | 0.254 (77) | 1.76 (10) | 7.19 (13) | 1.37 |
| 4 | 0.240 (72) | 1.41 (13) | 7.22 (15) | 1.49 |

### Polarization-gated fluorescence anisotropy

Polarization-gated fluorescence anisotropy of ANS in CHT was measured (Fig. 5) at different formalin concentrations to monitor changes around the ANS binding sites of the enzyme. The anisotropy decay in CHT (Figure 5; control) is characterized by three rotational time constants of ~ 56 ps (θ_1_), ~ 712 ps (θ_2_) and 30 ns (θ_3_), respectively, (Table 4). The faster (θ_1_) and the relatively slower correlation time (θ_2_) is attributed to orientational motion of the dye in water [11] and that (ANS) bound to an external binding site in CHT, respectively. On the other hand, the much longer time constant (θ_3_) originates from rotational motion of the ANS/CHT complex where the probe is strongly bound to an internal binding site of the enzyme so that it rotates with the enzyme as a protein/dye complex [23]. Upon interaction with formalin the fluorescence anisotropy decay of ANS in CHT is remarkably modified and a dip-rise pattern becomes evident (Fig 5) owing to heterogeneous distribution of ANS in different microenvironments with different fluorescence lifetimes and associated rotational correlation times [24–26]. When a fluorophore exists in different microenvironments, its anisotropy decay, r(t) can be expressed as: [27, 28]

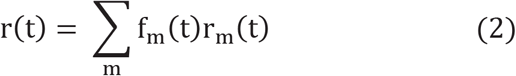

where the fractional intensity of the m^th^ component at any time t is given by

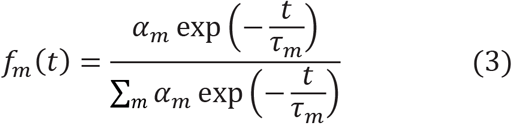

α_m_ is the amplitude corresponding to the fluorescence lifetime τ_m_ at t = 0 and

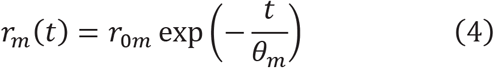

where θ_m_ is the single rotational correlation time of the m^th^ component and r_0m_ is corresponding initial anisotropy.

**Fig. 5:**
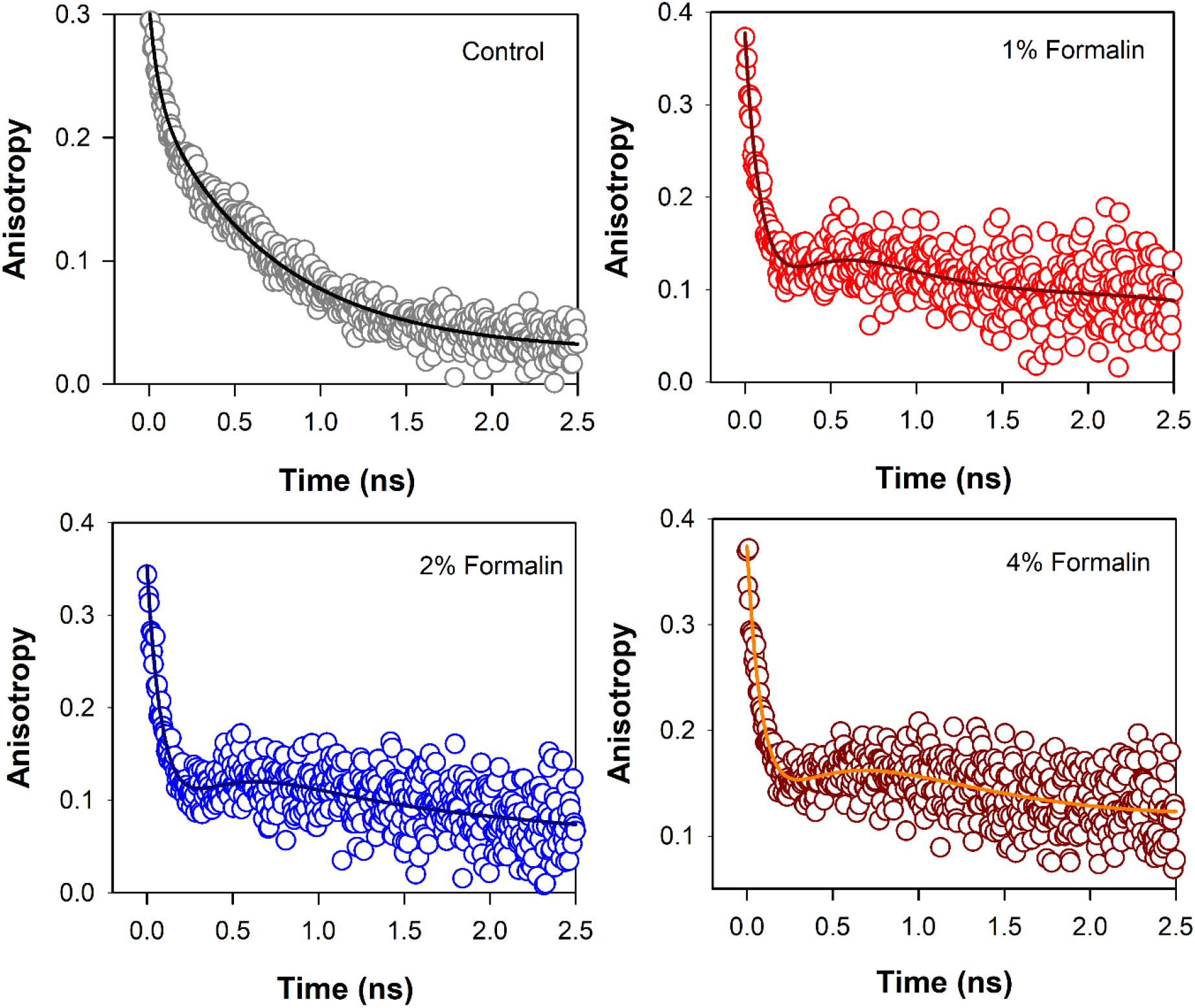
Anisotropy decay (r(t)) of ANS in CHT in absence and presence of formalin.

**Table. 4:** Time-resolved Fluorescence anisotropy parameters. The anisotropy decay of ANS in CHT is fitted by the following equation

| System | $\theta_1/\text{ns}$ | $\theta_2/\text{ns}$ | $\theta_3/\text{ns}$ |
| --- | --- | --- | --- |
| ANS in CHT | 0.056 | 0.712 | 30 |
| ANS in CHT+1 % formalin | 0.098 | 0.584 | 30 |
| ANS in CHT+2 % formalin | 0.094 | 0.425 | 30 |
| ANS in CHT+4 % formalin | 0.092 | 0.244 | 30 |

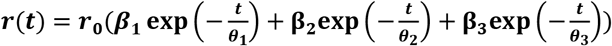

where r_0_ is initial anisotropy, β_is_ and θ_is_ are the amplitudes and corresponding rotational time constants, respectively. The longest rotational correlation time is kept constant at 30 ns. In presence of formalin, the anisotropy decays are fitted with equation (2) as described in the text.

The analysis of the dip and rise anisotropy decay (Fig. 5) using equation (2) shows presence of three rotational correlation times (Table 4) where the fastest (θ_1_) and the longest (θ_3_) rotational correlation times remain almost unchanged with a variation in the formalin content. However, a gradual decrease in the intermediate correlation time (θ_2_) from 712 ps to 244 ps with increasing formalin content is observed which may be attributed to covalent modification of the amino acid residues in the enzyme by formalin. As a result of such residue modification the external binding site of ANS in the enzyme likely becomes more exposed to water giving rise to faster rotational relaxation of the dye (ANS).

### Solvent accessible surface area (SASA)

Fig. 6 shows solvent accessible surface area of CHT (1CGJ). It becomes evident that several amino acid residues of CHT are exposed to solvent and hence, are vulnerable to formalin induced modification. However, we have restricted our study to the substrate and the probe (ANS) binding sites in the enzyme. At the catalytic S1 pocket of CHT which also serves as the buried internal site of ANS binding, Ser-195 is solvent exposed and contains a free hydroxyl group making it susceptible to formalin induced modification [7]. Furthermore, study of the external ANS binding site indicates Cys-1 to be vulnerable to modification by formalin owing to its high extent of solvent exposure and the presence of a free amino (-NH_2_) group. In contrary, Cys-122 has no free amino (NH_2_) group making it almost immune to formalin induced modification. This study also reveals the presence of susceptible residues Asn-48 and Ser-119 within 3Å of the ANS binding site, making them suitable candidates for our study.

**Fig. 6:**
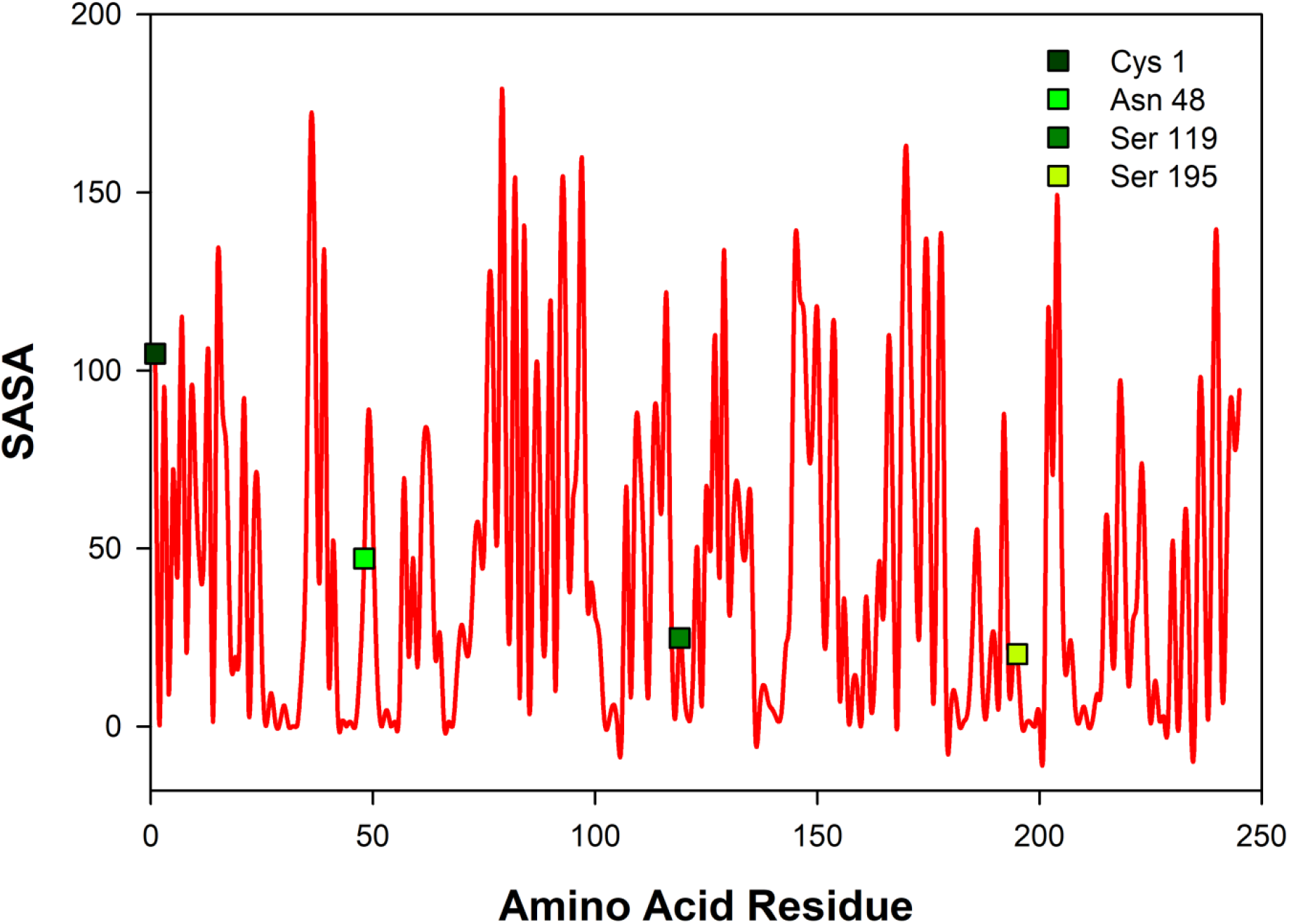
Solvent accessible surface area (SASA) of CHT [PDB ID: 1CGJ]. Four different residues corresponding to the catalytic site and molecular recognition were selected [Cys-1;Asn-48;Ser-119;Ser-195] for formalin induced modification.

### Residue modification and molecular docking analysis

To simulate and study the influence of minimal local residue modification on the function of CHT and its ability to recognize molecular binding partner, the residues selected from the SASA study (Ser195; Cys1; Asn48; Ser119) were subjected to residue modification according to Kamps et al. [7] Fig. 7 shows various possible modifications due to reaction with formalin. From Fig. 7 (a) it becomes evident that serine has three possible modified outcomes. The free hydroxyl (OH) group undergoes modification forming a methylol adduct (Ser’), which can undergo internal cyclization, forming oxazolidone adduct (Ser”) or forming a crosslink with the nearby His-57 residue (Ser’ “). Similarly, Cys-1 and Asn-48, undergo modifications forming a methylol adduct at the free thiol (-SH) or amino (-NH_2_) group (Fig. 7b & c). However, due to the formation of a disulfide bond between Cys-1 and Cys-122, no thiol group is available for formalin induced modification. Cys-1 being the N-terminal amino acid, contains a free amino group (-NH_2_) which reacts with the formalin.

**Fig. 7:**
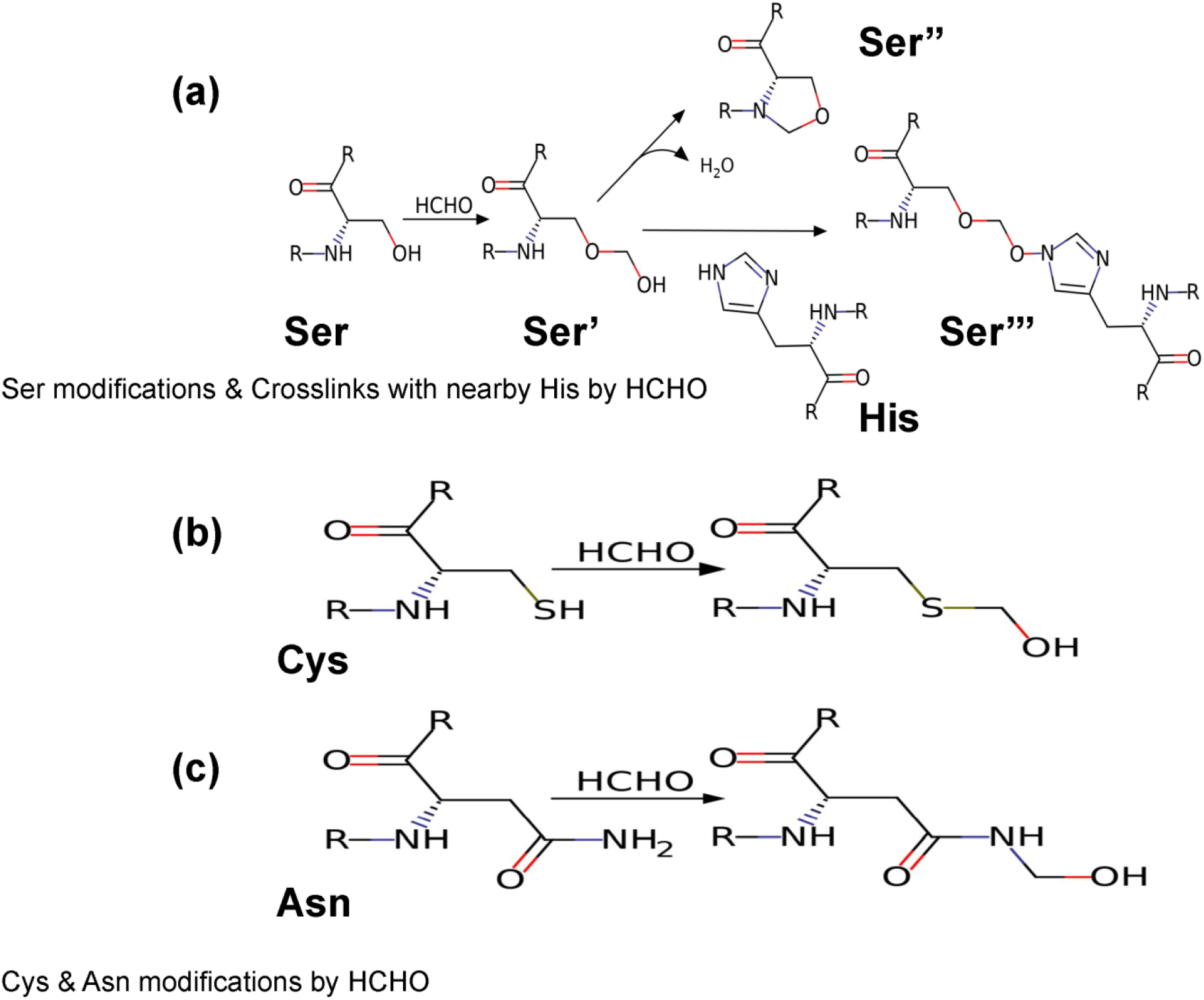
Reaction products of HCHO with different amino acids. (a) Amino acid Serine (Ser)reacting with HCHO form methylol derivative [Ser’]; which undergo cyclisation to form oxazolidine adduct [Ser’’]; the methylol derivative can react with nearby amino acid [case in point amino acid Histidine (His)] to form a crosslink derivative [Ser’’’]. (b & c) Reaction of HCHO with amino acids Cystine (Cys) and Asparagine (Asn) forming their respective methylol derivative.

Molecular docking was performed on the modified CHT structure. Fig. 8 & 9 & 10 illustrate different ligand bound structures. It becomes evident from the docking studies that residue modification at the catalytic site has no influence on binding of CHT to the substrate (AMC) (Fig. 8). Furthermore, a study of the van der Waals surface indicated no major structural distortions of the binding pocket (Supplementary Figure S.1). The possible formation of Ser-195-His-57 crosslink upon formalin treatment may cause a disruption of the serine protease activity at the catalytic triad; thus, impairing the product formation and release, leading to reduced catalytic activity.

**Fig. 8:**
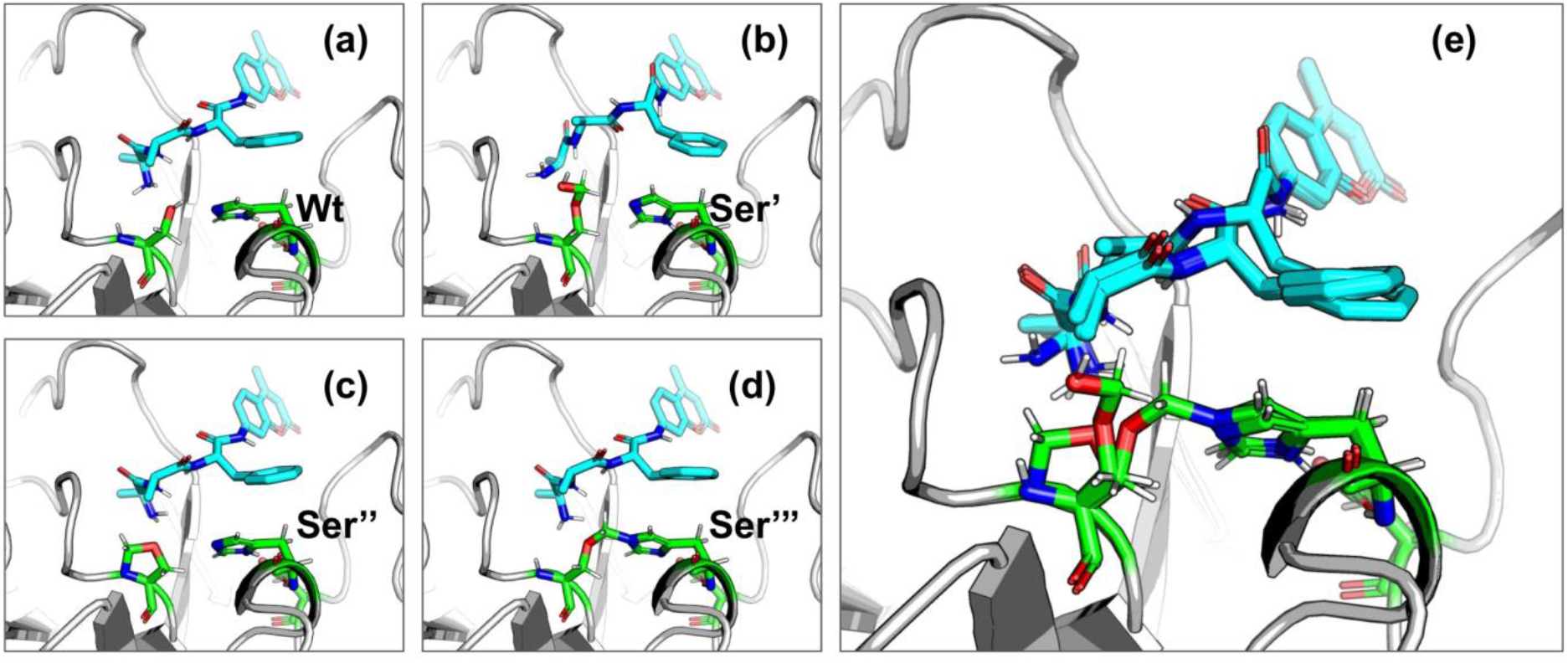
Binding of substrate (AMC) withWt and HCHO modified CHT as obtained by molecular docking simulations. (a-d) Different modes of AMC binding with CHT on modification of the active site Serine residue (Ser-195) as mentioned previously [Wt indicates the wild type Ser-195 residue]. (e) Overlapping view of the different binding interactions; residue modification causes no major change in binding mode of AMC with CHT.

**Fig. 9:**
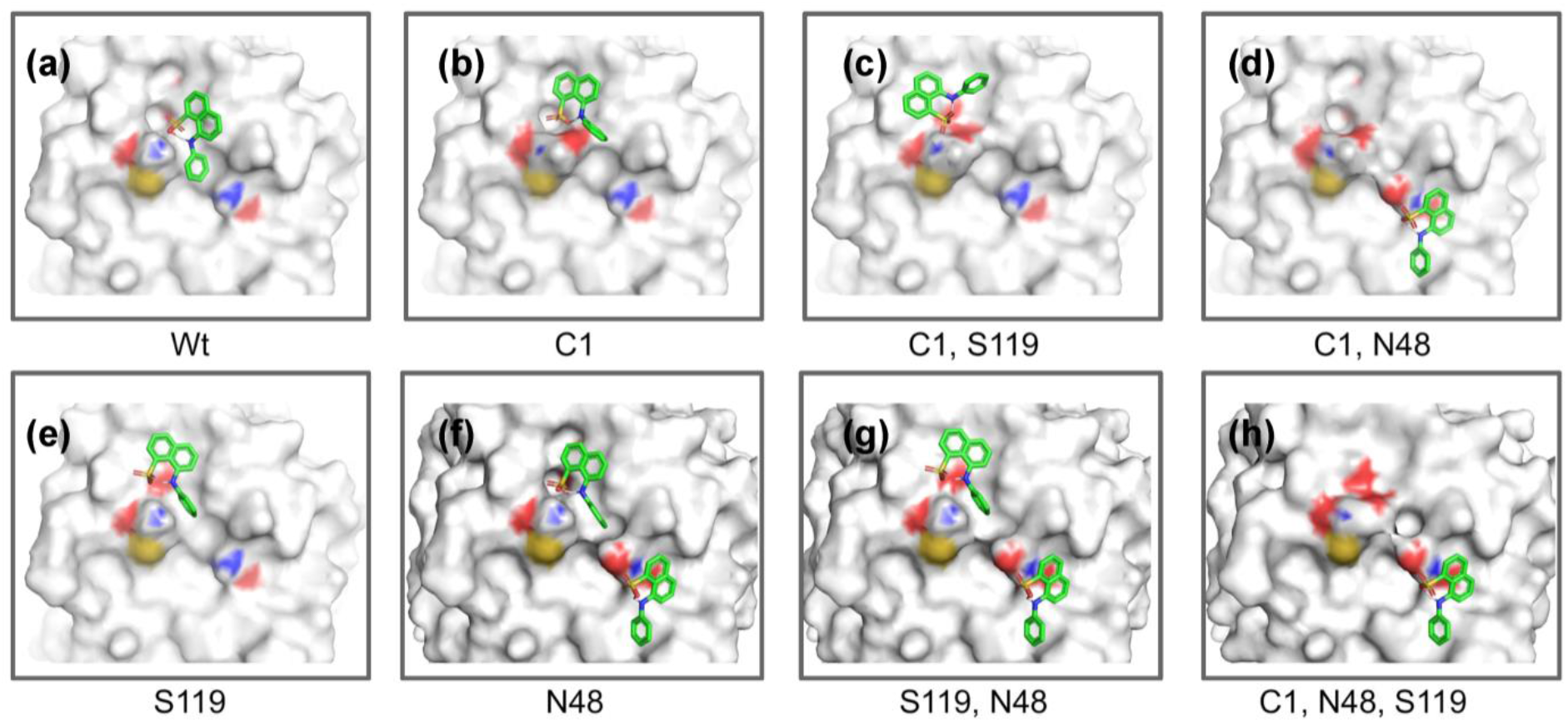
Different binding modes of ANS with CHT (external site) as obtained by molecular docking simulation. (a-h) Best binding modes of ANS with on residue modification [Cys-1;Asn-48;Ser-119] as mentioned previously; extensive change in binding modes resulting in heterogeneity of ANS binding is evident.

Molecular docking between ANS and CHT indicates presence of two binding sites, namely, the buried internal site at the S1 pocket and an external binding site at cysteine 1-122 di-sulfide bond. It becomes evident from (Fig. 9) that, formalin induced residue modification at the external binding site of ANS may lead to the generation of multiple probe (ANS) binding sites and a heterogeneous probe binding distribution which is indeed reflected by the dip-rise pattern in the anisotropy decay (Fig. 5). Docking simulations of CHT modified at the external binding site Cys-1 and its nearby susceptible residues Ser-119, Asn-48 generated binding models (Fig. 9b-h) which deviate from the unmodified CHT (Fig. 9a). Although molecular docking does not provide pronounced solvent effects in protein/ligand interactions, however, by probing the solvent accessibility of the bound ligand (ANS), solvent effects in protein-ligand interactions can be elucidated from molecular docking studies. SASA analysis of the bound probe found a diverse range of solvent effects upon formalin modification of the external site. The varied solvent effects ranged from mimicking the unmodified ANS binding site [modification at Cys-1;1244Å^2^ & Ser-119;1236Å^2^] and lesser degree of solvent accessibility [modifications at Asn-48;1150Å^2^] to experiencing higher solvent effects [modifications in conjunction at Cys-1 Asn-48;(1374Å^2^), Cys-1 Ser-119;(1402Å^2^), Ser-119 Asn-48;(1369Å^2^), Cys-1 Asn-48 Ser-119;(1360Å^2^)] (Table. 5a) signifying the bound probe to be more exposed and susceptible to solvent effects.

**Table. 5a:**
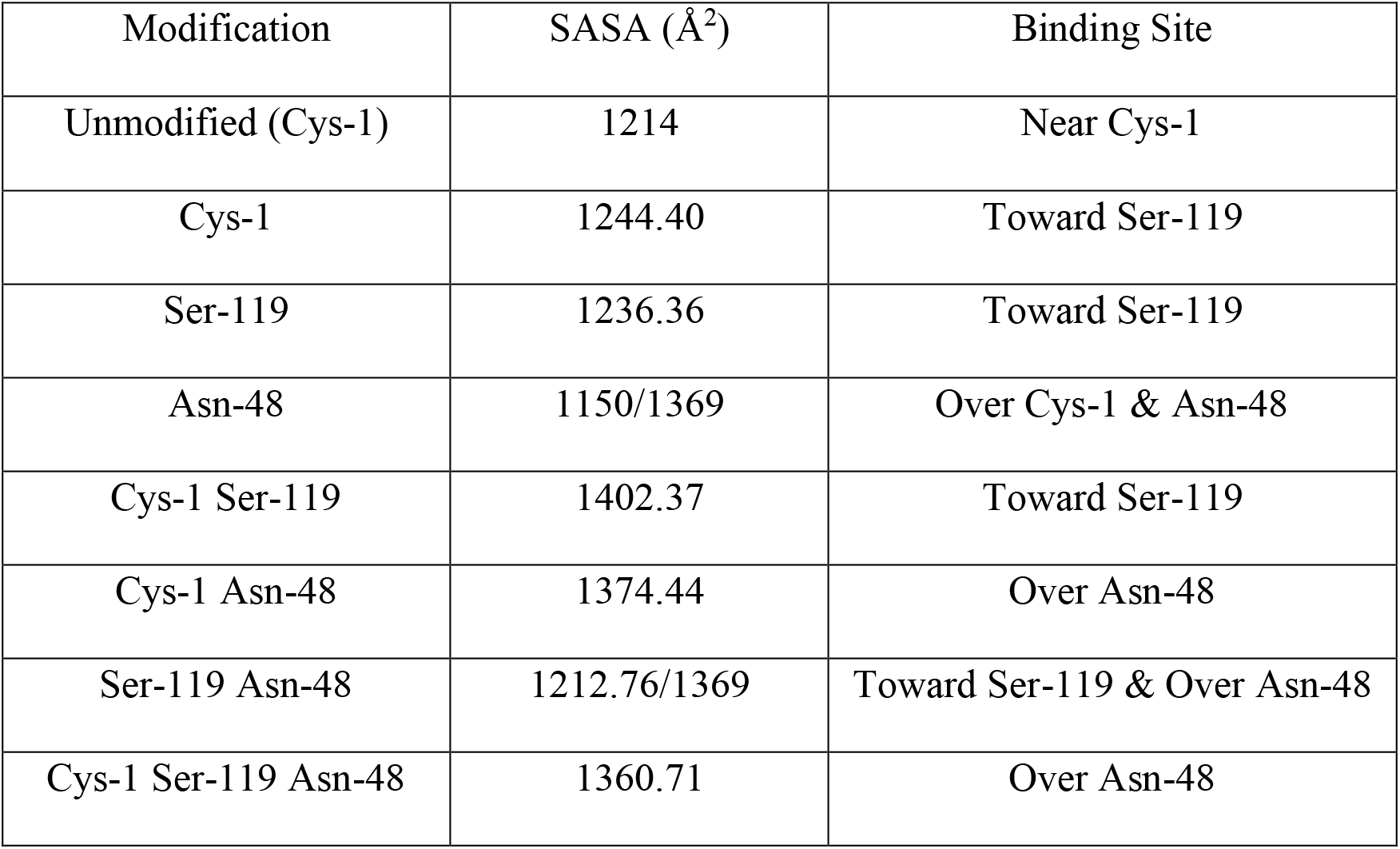
SASA of ANS upon binding to formalin modified CHT (External Site)

Similar to the binding interactions of substrate (AMC), residue modification at the internal ANS binding site in CHT (Ser-195) has no effect on binding of the probe (Fig. 10). SASA analysis of ANS at the internal binding site of CHT suggests no major changes in solvent exposure of ANS (Table. 5b) suggesting that the bound probe is shielded from the surrounding solvent even after residue modification.

**Fig. 10:**
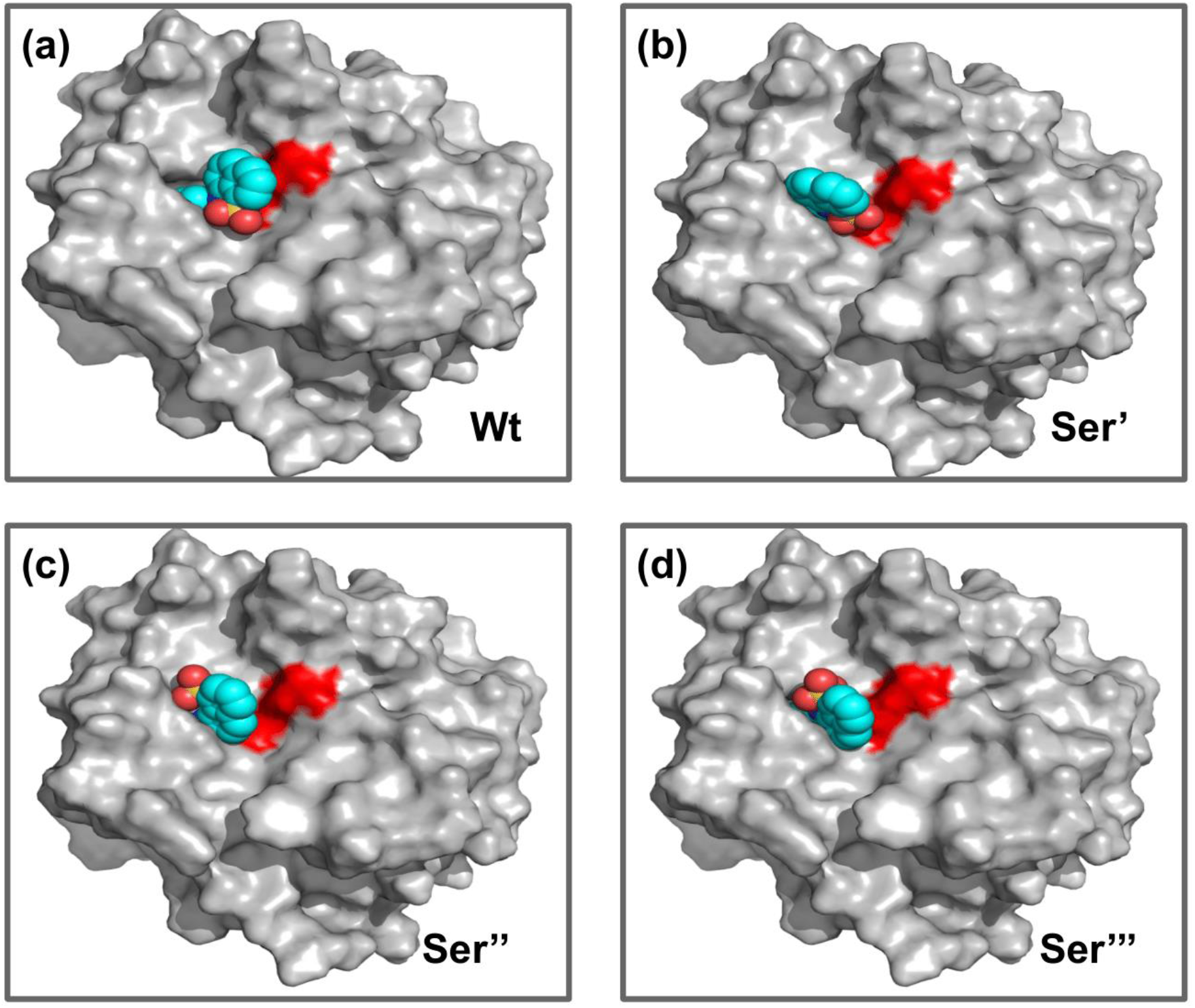
Different binding modes of ANS with CHT (internal site) as obtained by molecular docking simulation. (a-d) ANS binding modes with CHT on active site serine (Ser-195) modification as previously mentioned. [Wt indicates the wild type/unmodified CHT].

**Table. 5b:** SASA of ANS upon binding to formalin modified CHT (Internal Site)

| **Modifications** | **SASA (Å<sup>2</sup>)** |
| --- | --- |
| Unmodified Ser-195 | 1027 |
| Ser' | 1061 |
| Ser'' | 1024 |
| Ser''' | 1013 |

### Residue modification disrupts activity and introduces heterogeneity in molecular recognition

CHT causes the catalytic hydrolysis of AMC to 7-amido 4-methyl-coumarin.[29] The effect of formalin induced modification was monitored at different concentrations of formalin (1-4%). We have observed a gradual decrease in the rate of product formation with an increase in formalin concentration. However, stability of both the secondary and the tertiary structure indicated to local residue modification at the catalytic S1 pocket as the possible architect in reduction of catalytic activity of the enzyme in presence of formalin. Previous studies by Martin et al.[30] and Zhang et al.[31] found direct correlation between modification of the catalytic site residues and activity. Ser-195 at the catalytic triad was solvent exposed. Ser-195 modification and molecular docking showed no change in substrate (AMC) binding at the catalytic triad. Furthermore, no major structural distortions in the binding pocket were observed. The analysis implied a stable enzyme-substrate formation; however, disruption in the proton transfer cascade from Ser-195 to His-57 to Asp-102 due to serine-histidine cross-linking by formalin impairs product formation and release leading to the reduced catalytic activity. Molecular docking simulations further support that modifications in the local amino acid residues are solely responsible for the impaired catalysis.

Time resolved fluorescence studies of ANS in CHT manifest heterogeneity in binding of ANS upon formalin induced modification. As a result of formalin induced structural modification in the enzyme, ANS bound to an internal binding site becomes increasingly exposed to water thereby leading to a decreased fluorescence lifetime and rotational relaxation time.

Our SASA calculations identified four amino acid residues [Ser-195; Cys-1; Asn-48 & Ser-119] as one of the possible centers of local residue modification.

Molecular docking analysis of external ANS binding site indicated generation of new ANS binding sites having different solvent accessibility on formalin modification. However, no change in binding was observed for the internal binding site of ANS in CHT. The SASA of bound ANS suggested the bound probe to be buried and shielded from the surrounding solvent. Our solvent effect and docking simulation studies of the bound probe was found to corroborate with the time resolved fluorescence anisotropy studies manifesting heterogeneity in ANS binding to CHT following formalin induced modification.

### Conclusion

The present work primarily focuses on the influence of residue modification on the catalytic activity and molecular recognition of CHT. The study found remarkable changes in the catalytic activity and molecular recognition of the enzyme (CHT) upon formalin induced chemical modification of its amino acid residues. The structural and molecular docking analysis provide evidence of no impediment to the formation of a stable enzyme-substrate complex due to formalin induced modifications; rather impairment of the proton transfer cascade in the catalytic triad due to Ser-195-His-57 cross-linking leads to reduced catalytic activity. Picosecond resolved fluorescence studies reveal generation of multiple binding sites of a fluorophore ANS in CHT, and upon treatment with formalin a reduction in fluorescence lifetime and rotational relaxation time of the probe is observed when it is bound to an external binding site. Subsequent molecular docking studies and SASA analysis provide evidence towards heterogeneity of ligand (ANS) interaction with the enzyme (CHT) upon minimal change in the local amino acid residues. The findings of the present study offer a background towards fine tuning enzymatic capabilities, drug molecule recognition and protein engineering.

## ACKNOWLEDGMENT

SKP thanks the Indian National Academy of Engineering (INAE) for the Abdul Kalam Technology Innovation National Fellowship, INAE/121/AKF.

## Supplementary Information

### Supplementary Table

**Table. S1:**
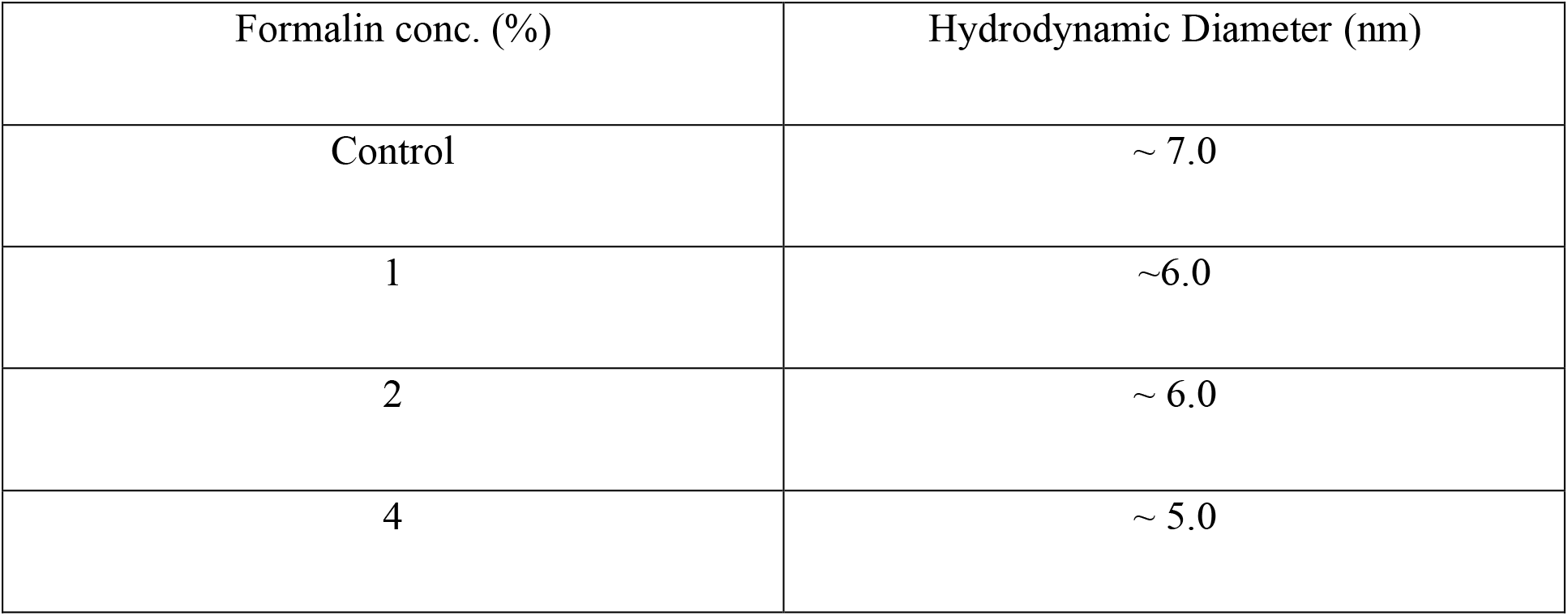
Globular Tertiary Structure of alpha-Chymotrypsin upon Formalin Treatment.

### Supplementary Figure

**Supplementary Fig.S1:**
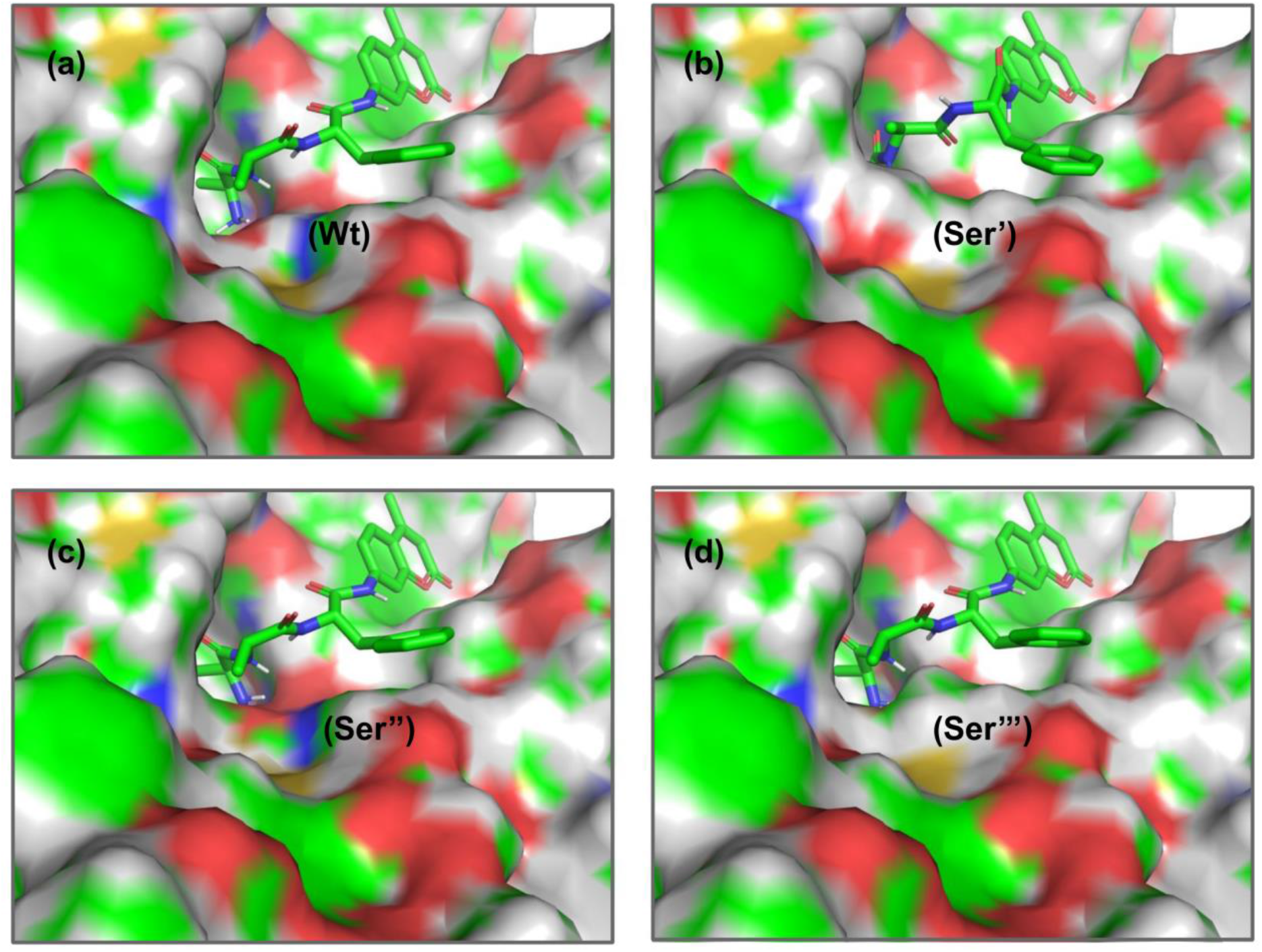
Surface view of the binding cavity at Wt and HCHO modified CHT. (a-d) Insignificant changes to the binding cavity upon residue modification at Ser-195.

## Reference

[1] H. Fraenkel-Conrat and H. S. Olcott, “Reaction of formaldehyde with proteins VI. crosslinking of amino groups with phenol, imidazole, or indole groups,” Journal of Biological Chemistry, vol. 174, pp. 827–843, 1948.

[2] E. A. Hoffman, B. L. Frey, L. M. Smith, and D. T. Auble, “Formaldehyde crosslinking: a tool for the study of chromatin complexes,” Journal of Biological Chemistry, vol. 290, pp. 26404–26411, 2015.

[3] D. Brutlag, C. Schlehuber, and J. Bonner, “Properties of formaldehyde-treated nucleohistone,” Biochemistry, vol. 8, pp. 3214–3218, 1969.

[4] B. Metz, G. F. Kersten, P. Hoogerhout, H. F. Brugghe, H. A. Timmermans, A. De Jong, et al., “Identification of formaldehyde-induced modifications in proteins reactions with model peptides,” Journal of Biological Chemistry, vol. 279, pp. 6235–6243, 2004.

[5] K. Lu, W. Ye, L. Zhou, L. B. Collins, X. Chen, A. Gold, et al., “Structural characterization of formaldehyde-induced cross-links between amino acids and deoxynucleosides and their oligomers,” Journal of the American Chemical Society, vol. 132, pp. 3388–3399, 2010.

[6] J. D. McGhee and P. H. Von Hippel, “Formaldehyde as a probe of DNA structure. 3. Equilibrium denaturation of DNA and synthetic polynucleotides,” Biochemistry, vol. 16, pp. 3267–3276, 1977.

[7] J. J. Kamps, R. J. Hopkinson, C. J. Schofield, and T. D. Claridge, “How formaldehyde reacts with amino acids,” Communications Chemistry, vol. 2, pp. 1–14, 2019.

[8] B. Metz, G. F. Kersten, G. J. Baart, A. de Jong, H. Meiring, J. ten Hove, et al., “Identification of formaldehyde-induced modifications in proteins: reactions with insulin,” Bioconjugate chemistry, vol. 17, pp. 815–822, 2006.

[9] B. Metz, T. Michiels, J. Uittenbogaard, M. Danial, W. Tilstra, H. D. Meiring, et al., “Identification of formaldehyde-induced modifications in diphtheria toxin,” Journal of Pharmaceutical Sciences, vol. 109, pp. 543–557, 2020.

[10] K. Lu, G. Boysen, L. Gao, L. B. Collins, and J. A. Swenberg, “Formaldehyde-induced histone modifications in vitro,” Chemical research in toxicology, vol. 21, pp. 1586–1593, 2008.

[11] S. K. Pal, J. Peon, and A. H. Zewail, “Ultrafast surface hydration dynamics and expression of protein functionality: α-Chymotrypsin,” Proceedings of the National Academy of Sciences, vol. 99, pp. 15297–15302, 2002.

[12] G. Böhm, R. Muhr, and R. Jaenicke, “Quantitative analysis of protein far UV circular dichroism spectra by neural networks,” Protein Engineering, Design and Selection, vol. 5, pp. 191–195, 1992.

[13] S. Sardar, S. Chaudhuri, P. Kar, S. Sarkar, P. Lemmens, and S. K. Pal, “Direct observation of key photoinduced dynamics in a potential nano-delivery vehicle of cancer drugs,” Physical Chemistry Chemical Physics, vol. 17, pp. 166–177, 2015.

[14] D. Bagchi, A. Ghosh, P. Singh, S. Dutta, N. Polley, I. I. Althagafi, et al., “Allosteric inhibitory molecular recognition of a photochromic dye by a digestive enzyme: dihydroindolizine makes α-chymotrypsin photo-responsive,” Scientific reports, vol. 6, pp. 1–11, 2016.

[15] G. F. Schröder, U. Alexiev, and H. Grubmüller, “Simulation of fluorescence anisotropy experiments: probing protein dynamics,” Biophysical journal, vol. 89, pp. 3757–3770, 2005.

[16] D. O’Connor, Time-correlated single photon counting: Academic Press, 2012.

[17] B. Chakraborty, C. Sengupta, U. Pal, and S. Basu, “Probing the hydrogen bond involving Acridone trapped in a hydrophobic biological nanocavity: Integrated spectroscopic and docking analyses,” Langmuir, vol. 36, pp. 1241–1251, 2020.

[18] O. Trott and A. J. Olson, “AutoDock Vina: improving the speed and accuracy of docking with a new scoring function, efficient optimization, and multithreading,” Journal of computational chemistry, vol. 31, pp. 455–461, 2010.

[19] W. Ma, C. Tang, and L. Lai, “Specificity of trypsin and chymotrypsin: loop-motion-controlled dynamic correlation as a determinant,” Biophysical journal, vol. 89, pp. 1183–1193, 2005.

[20] D. Banerjee and S. K. Pal, “Conformational dynamics at the active site of α-chymotrypsin and enzymatic activity,” Langmuir, vol. 24, pp. 8163–8168, 2008.

[21] A. K. Shaw, R. Sarkar, D. Banerjee, S. Hintschich, A. Monkman, and S. K. Pal, “Direct observation of protein residue solvation dynamics,” Journal of Photochemistry and Photobiology A: Chemistry, vol. 185, pp. 76–85, 2007.

[22] M. Collini, L. D’Alfonso, H. Molinari, L. Ragona, M. Catalano, and G. Baldini, “Competitive binding of fatty acids and the fluorescent probe 1-8-anilinonaphthalene sulfonate to bovine β-lactoglobulin,” Protein Science, vol. 12, pp. 1596–1603, 2003.

[23] J. D. Johnson, M. A. El-Bayoumi, L. D. Weber, and A. Tulinsky, “Interaction of. alpha.-chymotrypsin with the fluorescent probe 1-anilinonaphthalene-8-sulfonate in solution,” Biochemistry, vol. 18, pp. 1292–1296, 1979.

[24] J. R. Lakowicz, Principles of fluorescence spectroscopy: Springer science & business media, 2013.

[25] B. K. Paul and N. Guchhait, “Modulated photophysics of an ESIPT probe 1-hydroxy-2-naphthaldehyde within motionally restricted environments of liposome membranes having varying surface charges,” The Journal of Physical Chemistry B, vol. 114, pp. 12528–12540, 2010.

[26] S. S. Sinha, R. K. Mitra, and S. K. Pal, “Temperature-dependent simultaneous ligand binding in human serum albumin,” The Journal of Physical Chemistry B, vol. 112, pp. 4884–4891, 2008.

[27] J. Broos, A. J. Visser, J. F. Engbersen, W. Verboom, A. van Hoek, and D. N. Reinhoudt, “Flexibility of enzymes suspended in organic solvents probed by time-resolved fluorescence anisotropy. Evidence that enzyme activity and enantioselectivity are directly related to enzyme flexibility,” Journal of the American Chemical Society, vol. 117, pp. 12657–12663, 1995.

[28] B. K. Paul, N. Ghosh, and S. Mukherjee, “Interplay of multiple interaction forces: binding of norfloxacin to human serum albumin,” The Journal of Physical Chemistry B, vol. 119, pp. 13093–13102, 2015.

[29] R. Biswas and S. K. Pal, “Caging enzyme function: α-chymotrypsin in reverse micelle,” Chemical physics letters, vol. 387, pp. 221–226, 2004.

[30] C. J. Martin, N. B. Oza, and M. A. Marini, “Formaldehyde as an Active Site Label of α-Chymotrypsin,” Canadian Journal of Biochemistry, vol. 50, pp. 1114–1121, 1972.

[31] W.-N. Zhang, D.-P. Bai, X.-Y. Lin, Q.-X. Chen, X.-H. Huang, and Y.-F. Huang, “Inactivation kinetics of formaldehyde on N-acetyl-β-d-glucosaminidase from Nile tilapia (Oreochromis niloticus),” Fish physiology and biochemistry, vol. 40, pp. 561–569, 2014.

